## Supplementary Information for "Theory of mechano-chemical patterning in biphasic biological tissues"

### 1 Results

#### 1.1 Derivation of the model

##### 1.1.1 Tissue spatial organisation

As sketched in Fig. 1 of main text, we model multicellular tissues as biphasic porous media, with a first phase consisting in a three-dimensional elastic meshwork made of interconnected cells, and a second phase being made up of aqueous extracellular fluid permeating in-between cells. The typical lengthscale of the whole tissue is $l$. Cells, compartmentalized by their plasma membrane have a typical diameter $l_c$ and the average intercellular or interstitial space has a characteristic size $l_i$. Typically, these three lengthscales are well-separated such that:

$$l \gg l_c \gg l_i.$$

We adopt a continuum description of the tissue where all physical quantities are averaged over a representative volume element (RVE) of intermediate size $l \gg l_r \gg l_c$ at which the system can be coarse-grained [1, 2]. At that scale, we introduce the space coordinate $\vec{r} \in \Omega$, where $\Omega$ is the domain occupied by the whole tissue, and $t$ denotes the time coordinate.

##### 1.1.2 Derivation of the biphasic model

Although we only consider a biphasic media in the main text of this manuscript, a more complete version of the model would introduce three volume fractions within the tissue: $\phi_s(\vec{r}, t)$ the volume occupied by the skeletal meshwork of cells divided by the volume of the RVE, $\phi_c(\vec{r}, t)$ the volume of fluid contained within cell membranes divided by the volume of the RVE and $\phi_e(\vec{r}, t)$ the volume occupied by interstitial fluid divided again by the RVE. In the following, we provide a rationale and orders of magnitude for considering only two phases in the main text.

As a consequence of volume conservation, we have here:

$$\phi_s + \phi_c + \phi_e = 1.$$

A difference with the biphasic model is that there are now explicitly three velocities associated with each of these phases, $v_s$, $v_c$ and $v_e$. Importantly, although it could be thought that fluid is only exchanged between cells and extracellular medium (which would have meant that $v_c = v_s$, i.e. solid phase and intracellular fluid are advected together), this is not the case, as gap junctions allow for direct cell-cell communication and fluid exchange (which in the three phase model mean that there can be non-zero $v_c$ with vanishing $v_s$). This is important when writing intracellular morphogen concentration laws, as $v_s$ would be included in an advective term, whereas $v_c$ would not, as most morphogens are too large molecules to be transported through gap junctions. In order to simplify the description of the system, we make two important assumptions relative to the cellular meshwork. First, as both cells and extracellular space are mostly composed of fluid, we neglect the volume fraction occupied by the skeletal meshwork of cells such that $\phi_c \stackrel{\text{def}}{=} \phi$ and $\phi_e \simeq 1-\phi$. Then, we assume that the velocity of cells moving in the tissue is very slow compared to the velocity of the fluid flows in the extracellular space and from cell to cell. Thus, the present study focuses on cell specification patterns considering that the evolution of the tissue topology is quasi-static. To make this approximation clearer, we provide below (Section 1.1.3) simple physical estimates showing that the characteristic timescales associated with tissue remodelling processes, such as cell proliferation and active cell motility, can be larger by an order of magnitude than fluid circulation inside the tissue.

In Section. 1.9, we also show how the theory can be simply extended to account for large scale cellular remodelling occurring during embryonic development and changing the tissue topology on a timescale similar to that of the fluid flows inside the tissue.

##### 1.1.3 Fluid re-circulation vs tissue remodelling

Large scale advective flows of extracellular fluid must be compensated by some recirculation of material ensuring global volume conservation (except if there are exchanges with other tissues, which we do not consider here for the sake of simplicity). We first consider the possibility of this re-circulation to occur through intracellular flows, and calculate the time scales associated with this. In general, there are three characteristic timescales associated with fluid circulation inside a multicellular tissue: $\tau_i$ for fluid flows inside the interstitial space, $\tau_c$ for fluid flows from one cell to the other through gap junctions, and finally $\tau$ for the exchanges of fluid between cells and the extracellular fluid through aquaporins.

As we demonstrate in Section 1.1.7 using a poroelastic model of multicellular tissues, $\tau_i$ and $\tau_c$ can be interpreted

*LIPHY, CNRS UMR 5588 & Université Grenoble Alpes, F-38000 Grenoble, France.

†Cavendish Laboratory, Department of Physics, University of Cmbridge, Cambridge, CB3 0HE, UK; Wellcome Trust/CRUK Gurdon Institute, University of Cambridge, Cambridge, CB2 1QN, UK; Wellcome Trust/MRC Stem Cell Institute, University of Cambridge, Cambridge, CB2 1QR, UK.

‡IST Austria, Am Campus 1, 3400 Klosterneuburg, Austria.

$$\tau_i = \eta l_p^2/(\kappa K) \text{ and } \tau_c = \eta l_p^2/(\kappa_c K)$$

where $\eta \approx 10^{-3}$Pa.s is the viscosity of the aqueous cytosolic fluid [3, 4], $K \approx 10^4$ Pa is the drained elastic modulus of the tissue [5], $l_p \approx 10^{-4}$ m the characteristic size of multicellular patterns in the tissue. $\kappa$ and $\kappa_c$ (expressed in [m$^2$]) are the effective interstitial and cell to cell water permeabilities. In agreement with the typical timescales involved in cell volume changes following an osmotic perturbation [6], we estimate that $\tau \approx 10^2 - 10^3$s. In the main text, based on typical tissue interstitial sizes and packing fractions, we estimate the interstitial permeability $\kappa \approx 10^{-15} - 10^{-19}$ m$^2$ leading to $\tau_i \approx 1 - 10^4$ s. We note that this estimate is one of the coarsest one in the model, as the packing fraction of multicellular tissues can vary widely between tissues and species.

Finally, the cell-cell permeability relies on gap junctions which are highly specialized membrane structures made of clusters of transmembrane protein channels named connexons and which connect together the cytoplasms of neighbouring cells [7]. The ensuing permeability can be estimated in applying the Darcy-Weisbach equation to a parallel array of connexons mediating the cell to cell fluid flux:

$$\kappa_c = \frac{d^2}{32}\left(1 + n\frac{d^2}{\delta l_c}\right) \approx 3 \times 10^{-18}\,\text{m}^2$$

taking, $l_c \approx 10^{-5}$ m the diameter of a cell, $n \approx 10^5$ as the average number of connexons at a cell-cell interface [8–11], $d \approx 3 \times 10^{-9}$ m [7] as the typical diameter of a connexon and $\delta \approx 10^{-8}$ m as its length [7, 12]. The coefficient 32 is the friction factor for a laminar flow [13]. Interestingly, this simple theoretical estimate of $\kappa_c$ is in line with direct experimental measurements using dyes in epithelial cell monolayers [14, 15]. Eventually, this provides an estimate of $\tau_c \approx 4 \times 10^2$ s.

Next, we compare these timescales of direct cell-cell fluid re-circulation to timescale of tissue remodelling. Indeed, an alternative to enforce global conservation given extracellular fluid flows would be to have a counter-flow of the celular meshwork. Typically, this is very slow, as the viscosity of multicellular tissues is very high. Such timescale have been estimated from rheological measurements made on various multicellular tissues [16, 17] and are typically $\tau_r \approx 10^4$s, associated with slow tissue re-arrangement and remodelling. These slow timescales are consistent with the observed timescales for cell division and cell motility, which have been both noted as major sources of tissue remodelling [18]. The typical timescale of cell division in embryonic tissues is $\tau_d \approx 10^4 - 10^5$s. The characteristic velocity of a cell moving within such a tissue is $v_m \approx 10^{-9}$m.s$^{-1}$. Over the characteristic size $l_p \approx 10^{-4}$ m of tissue patterning, this leads to a typical timescale for motility-driven remodelling of $\tau_m \approx 10^5$s.

Importantly, we see that all three timescales associated with fluid flows are either slightly or much smaller compared to the timescale associated with tissue remodelling and cellular flows. Therefore, in our model, we consider tissue remodelling as a quasi-static process and assume a fixed tissue structure during the faster fluid exchanges between cells or between cells and extracellular fluid. A more complete description incorporating these effects and wich does not make such a simplifying assumption is presented in Section 1.9. This is motivated by the fact that in some biological systems, during early embryogenesis, cell division could be faster than our estimate, closer to $\tau_d \approx 10^3$s. In this situation, plastic re-arrangements could be potentially more important and play a role in the patterning instabilities we describe.

###### 1.1.4 Fluid conservation

As fluid can be exchanged between cells and between cells and extracellular medium, we can write, assuming fluid incompressibility, the following mass balance relations:

$$\begin{aligned} \partial_t \phi + \nabla.(\phi v_c) &= S \\ \partial_t (1-\phi) + \nabla.((1-\phi) v_e) &= -S \end{aligned} \quad (1)$$

where $v_{c,e}(\vec{r}, t)$ are velocities associated with fluid flows respectively inside and outside the cells. It should be noted that because these equations are defined at a coarse-grained level, $v_c$ refers to fluid flowing directly from one cell to the other, which occurs in multicellular tissues via gap junctions, as discussed in the previous section. Gap junctions have been well-characterised as key regulators of cell-cell coupling, allowing in addition of fluid transport, the movement of inorganic ions and small water-soluble molecules from the cytoplasm of one cell to the other [19]. Nevertheless, most morphogens cannot cross through gap junctions, as they are proteins whose size exceeds by far the diameter of connexons lumen [7, 12]. Crucially, in our model, this means that although extracellular flows $v_e$ can advect morphogens, this is not the case of intercellular flows $v_c$. This allows advection to concentrate morphogens locally, as they do not participate in fluid re-circulation.

Summing up the two previous mass balance equations, we obtain $\nabla.(\phi v_c + (1-\phi) v_e) = 0$ which is generally satisfied if:

$$v_c = -\frac{1-\phi}{\phi} v_e.$$

As $\phi$ is usually rather close to one, the cell to cell fluid velocity $v_c$ is thus expected to be much smaller than the velocity of fluid flows developing in the extracellular space in-between cells $v_e$. However, in spite of taking relatively low values, $v_c$ should not be neglected, as it is central for the model consistency and in enforcing volume conservation in the system. We note here that although the cell volume fraction $\phi$ is close to one in many tightly packed multicellular tissues, our theory does not depend on that assumption, and can in principle accommodate values of $\phi$ very different from one.

The source term $S$, on the right-hand side of Eq. 1, originates both in the genetically controlled transport of water between cells and extracellular medium, which occurs through specific transmembrane proteins named aquaporins [20, 21] and on the balance between cell division and apoptosis regulating the nuber of cells in a RVE. In absence of spatial fluid flows in interstitial spaces or at tissue boundaries, we have:

$$S = \partial_t \phi = \partial_t (N_c V_c)/V_r$$

where $N_c$ is the number of cells in a RVE, $V_c$ the volume of a single cell and $V_r$ the volume of an RVE. When submitted to an external osmotic shock or mechanical load, the volume of cells transiently increases or decreases prior to relaxing to an homeostatic volume $V_h$. Considering an exponential relaxation, we can write:

$$\partial_t V_c = \frac{V_h - V_c}{\tau}$$

where $\tau$ is the typical timescale associated with a regulatory response to cell volume changes. This time scale can be estimated to be of the order of $\sim 10^2 - 10^3$ s in the case of an osmotic shock [6].

In our model, as discussed with the orders of magnitude provided in Section 1.1.3, the tissue topology is taken to be fixed and we thus neglected the variations of $N_c$ due to cell division and cell loss. However, we present below a more general argument and show that these phenomena does not impact the expression of the term $S$ in (1). This argument will be also used to justify the general model presented in Section 1.9 where tissue topology becomes dynamic thanks to cell motility and cell turnover.

The simplest evolution law for $N_c$ is given by $\partial_t N_c = s_d - N_c k_a$, where $s_d$ accounts for cell division and $k_a$ is the cell loss rate. This law can be rewritten as

$$\partial_t N_c = \frac{N_h - N_c}{\tau_d},$$

where $\tau_d = k_a^{-1}$ is the typical duration of a cell cycle of the order of $\sim 10^4 - 10^5$ s (during chick limb bud patterning this time is about 10 $h = 3.6 \times 10^4$ s) and $N_h = s_d \tau_d$ is the equilibrium cell number when cell division balances cell loss. Defining $\phi_h = N_h V_h / V_r$ the homeostatic volume fraction of cells and $\delta V$ and $\delta N$ some initial perturbations from the homeostatic state, we obtain,

$$S = \frac{\tau + \tau_d}{\tau \tau_d}(\phi_h - \phi) + \phi_h \left( \frac{\delta V}{V_h} \tau_d^{-1} \mathrm{e}^{-t/\tau} + \frac{\delta N}{N_h} \tau^{-1} \mathrm{e}^{-t/\tau_d} \right).$$

Considering typical timescales such that $t \gg \tau$ and as $\tau_d \gg \tau$, the first term dominates the last expression and we finally obtain $S = (\phi_h - \phi)/\tau$.

In agreement with the fact that cell fate decisions are usually accompanied by modifications of the cell cycle and cell morphology, including cell volume changes, we suppose that $\phi_h$ depends on the concentrations of morphogens [22–24]. Indeed, most of the known morphogens are in fact growth factors which regulate both cell differentiation but also cell growth and cell division [25–28]. We can thus write the source term of the conservation equation as:

$$S = \frac{\phi_h(A_i, I_i) - \phi}{\tau}$$

where $\phi_h(A_i, I_i)$ is the cell volume fraction with cells in their homeostatic volume state.

As mentioned above, it should be noted for the sake of completeness that we have assumed here a closed-system, where no fluid is exchanged with the rest of the organism. Relaxing this assumption would be equivalent to adding another source term, $S_b$, in the conservation equation for $1 - \phi$ (see Eq. 1), without adding it to the conservation equation for $\phi$. This would be an alternative to intracellular re-circulation to allow for the existence of non-zero steady state fluid flows in the tissue.

###### 1.1.5 Morphogen conservation outside cells

Conservation equations for morphogens contained in the extracellular space are given by:

$$\begin{aligned} \partial_t((1-\phi)A_e) + \nabla.\left(A_e(1-\phi)v_e - D\nabla A_e\right) &= -\gamma_A A_e + \lambda_A A_i \\ \partial_t((1-\phi)I_e) + \nabla.\left(I_e(1-\phi)v_e - D\nabla I_e\right) &= -\gamma_I I_e + \lambda_I I_i \end{aligned} \quad (2)$$

where $\gamma_{A,I}$ and $\lambda_{A,I}$ are respectively the import and export rates of morphogens through some specific channels and pumps, and $D$ is the common global diffusion coefficient of both morphogens in the intercellular space. $D$ do not refer to a molecular diffusion coefficient but to an homogenized quantity over a RVE.

###### 1.1.6 Morphogen conservation inside cells

Conservation equations for morphogens inside cells read:

$$\begin{aligned} \partial_t(\phi A_i) &= f(A_i, I_i) + \gamma_A A_e - \lambda_A A_i \\ \partial_t(\phi I_i) &= g(A_i, I_i) + \gamma_I I_e - \lambda_I I_i \end{aligned} \quad (3)$$

where $f$ and $g$ are the morphogens turnover rates, non-linear functions of morphogen concentrations describing their production and degradation by cells. Note here, that these reaction terms could also be a function of extracellular morphogen concentrations, describing a biological situation where signaling at the membrane also directs internal morphogen production and degradation but that would not impact our final model (10) as explained in Section 1.3.

The vector field $[f, g]$ is assumed to have a single stable equilibrium point $(A_i^*, I_i^*)$ such that $f(A_i^*, I_i^*) = g(A_i^*, I_i^*) = 0$ and det $\mathbb{J} > 0$ and tr $\mathbb{J} < 0$ such that the eigenvalues of the jacobian matrix

$$\mathbb{J} = \begin{pmatrix} \partial_{A_i^*} f & \partial_{I_i^*} f \\ \partial_{A_i^*} g & \partial_{I_i^*} g \end{pmatrix}$$

are negative and we do not consider the complex case where morghogens can undergo transitions between various stable chemical states.

Contrary to (2), advective terms are absent in (3), as known morphogens are too large to be transported through gap junctions [7, 12]. Indeed, because of this impossibility of direct cell to cell transport of morphogens, advective terms would equate the transport terms $\nabla(\phi A_i v_s)$ and $\nabla(\phi I_i v_s)$, where $v_s(\vec{r}, t)$ is the cellular meshwork velocity, which we neglect here by ignoring cellular motions. Such assumption is critically discussed in Section 1.9.

###### 1.1.7 Effective mechanical behaviour of the tissue

To complete the model described by Eqs. (1,2,3), we need to relate the interstitial fluid velocity, $v_e$, with the cell volume fraction $\phi$. Such link is modelled using the classical poroelastic theory [29, 30].

When biochemical reactions involving morphogens are at equilibrium, (1) has an homogeneous solution given by:

$$v_e = 0 \text{ and } \phi \overset{\text{def}}{=} \phi^* = \phi_h(A_i^*, I_i^*)$$

This state of the tissue constitutes a "reference configuration" where the deformation of the multicellular network is considered to be null. Supposing small variations from this reference state, we can write an effective quadratic mechanical free energy for the cellular network, $F$, coupling the tensorial infinitesimal strain $\mathbb{E}$ with porosity variations $\delta\phi = \phi - \phi^*$. The free energy of a poroelastic tissue reads:

$$F = G\mathbb{E} : \mathbb{E} + \left(\frac{K_u}{2} - \frac{G}{3}\right)[\text{tr }(\mathbb{E})]^2 + \frac{K_u - K}{\alpha}\text{tr }(\mathbb{E})\delta\phi + \frac{K_u - K}{2\alpha^2}[\delta\phi]^2$$

where $G$ is the shear modulus of the tissue, $K$ is its drained modulus, $K_u$ its undrained modulus and $\alpha$ its Biot coefficient.

We derive the non-dissipative mechanical stress in the tissue, $\sigma$, and the interstitial fluid pressure $p$ by differentiation of the free energy:

$$\begin{array}{c} \sigma = \partial_\epsilon F = 2G\mathbb{E} + \left(K - \frac{2G}{3}\right)\text{tr}(\mathbb{E})\mathbb{I} - \alpha p\mathbb{I} \\ p = -\partial_{\delta\phi}F = -\frac{K_u - K}{\alpha}\text{tr}(\mathbb{E}) - \frac{K_u - K}{\alpha^2}\delta\phi, \end{array} \tag{4}$$

where $\mathbb{I}$ denotes the identity matrix. In this classical theory, mechanical energy dissipation is assumed to happen mainly through viscous friction of the extracellular fluid permeating in between the cells and is described using Darcy's law:

$$(1 - \phi)v_e = -\frac{\kappa}{\eta}\nabla p,$$

as, again, we neglect $v_s$ in this framework, but this time as compared to the extracellular fluid velocity $v_e$. The tissue permeability is denoted $\kappa$ ([m$^2$]) and $\eta$ is the viscosity of the permeating aqueous fluid.

Finally, force balance inside the tissue implies that $\nabla.\sigma = 0$, which using the first constitutive relation (4) leads to

$$2G\nabla.\mathbb{E} + \left(K - \frac{2G}{3}\right)\nabla\text{tr}(\mathbb{E}) = \alpha\nabla p.$$

Coupling that last equation with the second equation of (4) and assuming, in line with the typical orders of magnitude for biological tissues [31, 32] that

$$K_u \gg K \gg G \text{ and } \alpha \simeq 1,$$

we obtain the simple relation:

$$(1 - \phi)v_e = -\frac{\kappa}{\eta}\nabla p = \frac{K\kappa}{\eta}\nabla\phi, \tag{5}$$

relating $v_e$ and $\phi$ where we can introduce the hydrodynamic diffusion coefficient of extracellular fluid through the tissue $D_m = K\kappa/\eta$, which is, as seen in the main text, a driving force of the instability, counter-acting morphogen Fickian diffusion which acts as a resisting force.

#### 1.2 Model of an active biphasic tissue

Coupling Eqs. (1,2,3,5), the full set of equations describing the chemo-mechanical behaviour of an active biphasic multicellular tissue can be written as:

$$\begin{array}{c} \partial_t(\phi A_i) = f(A_i, I_i) + \gamma_A A_e - \lambda_A A_i \\ \partial_t(\phi I_i) = g(A_i, I_i) + \gamma_I I_e - \lambda_I I_i \\ \partial_t\left((1 - \phi)A_e\right) + \nabla.\left(A_e D_m \nabla\phi - D\nabla A_e\right) = -\gamma_A A_e + \lambda_A A_i \\ \partial_t\left((1 - \phi)I_e\right) + \nabla.\left(I_e D_m \nabla\phi - D\nabla I_e\right) = -\gamma_I I_e + \lambda_I I_i \\ \partial_t\phi - \nabla.(D_m\nabla\phi) = \frac{\phi_h(A_i, I_i) - \phi}{\tau}. \end{array}$$

This system of coupled equations can also be written in a more compact matrix form:

$$\begin{array}{c} \partial_t(\phi \underline{X}_i) = \underline{F}(\underline{X}_i) + \mathbb{\Gamma}\underline{X}_e - \mathbb{\Lambda}\underline{X}_i \\ \partial_t((1 - \phi)\underline{X}_e) + \nabla.\left(D_m\nabla\phi\underline{X}_e - D\nabla\underline{X}_e\right) = -\mathbb{\Gamma}\underline{X}_e + \mathbb{\Lambda}\underline{X}_i \\ \partial_t\phi - \nabla.(D_m\nabla\phi) = \frac{\phi_h(\underline{X}_i) - \phi}{\tau}, \end{array} \tag{6}$$

where $\underline{X}_{i,e} = (A_{i,e}, I_{i,e})$ are the morphogen concentration vectors, $\underline{F}(\underline{X}_i) = (f(A_i, I_i), g(A_i, I_i))$ is the morphogen turnover rates vector, and

$$\mathbb{\Lambda} = \begin{pmatrix} \lambda_A & 0 \\ 0 & \lambda_I \end{pmatrix} \text{ and } \mathbb{\Gamma} = \begin{pmatrix} \gamma_A & 0 \\ 0 & \gamma_I \end{pmatrix}$$

are the diagonal matrices of transmembrane import and export rates of morphogens.

The model has as a single homogeneous steady state solution given by:

$$\phi \equiv \phi^*, \ \underline{X}_i = \underline{X}_i^* \text{ and } \underline{X}_e = \mathbb{K}\underline{X}_i^* \tag{7}$$

where $\mathbb{K} = \mathbb{\Lambda}\mathbb{\Gamma}^{-1}$ is the matrix of the transmembrane transport equilibrium constants.

To investigate the stability of this solution and fully specify the problem, we need, in addition of (6), to define initial and boundary conditions. The initial condition is typically chosen as a random perturbation of the homogeneous solution (7). For the boundary conditions on the frontier $\partial\Omega$ of the domain $\Omega$, we impose a no-flux of fluid and morphogens:

$$\nabla\underline{X}_e.n|_{\partial\Omega} = \nabla I_e.n|_{\partial\Omega} = \nabla\phi.n|_{\partial\Omega} = 0,$$

where $n$ is the outward normal of $\Omega$. Such conditions are natural as they correspond to an isolated system and we wish to consider endogenous pattern formation that are not driven externally.

Eventually, we also need to specify the non-linear function $f, g$ and $\phi_h$ as a function of the internal morphogens concentrations. For $f$ and $g$, we will essentially consider two cases in this paper:

For Fig. 2 of the main text, we use the classical activator-inhibitor Gierer-Meinhardt scheme [33–35] (see section "Turing-Keller-Segel instabilities" in the main text):

$$\begin{array}{c} f(A_i, I_i) = \rho\frac{A_i^2}{I_i} - \frac{A_i}{\tau_A} \\ g(A_i, I_i) = \rho A_i^2 - \frac{I_i}{\tau_I}, \end{array} \tag{8}$$

where $\rho$ is the rate of activation and inhibition and $\tau_{A,I}$ the timescales of degradation of $A$ and $I$. Within this scheme, $A_i^* = \tau_A/\tau_I$ and $I_i^* = \rho\tau_A^2/\tau_I$.

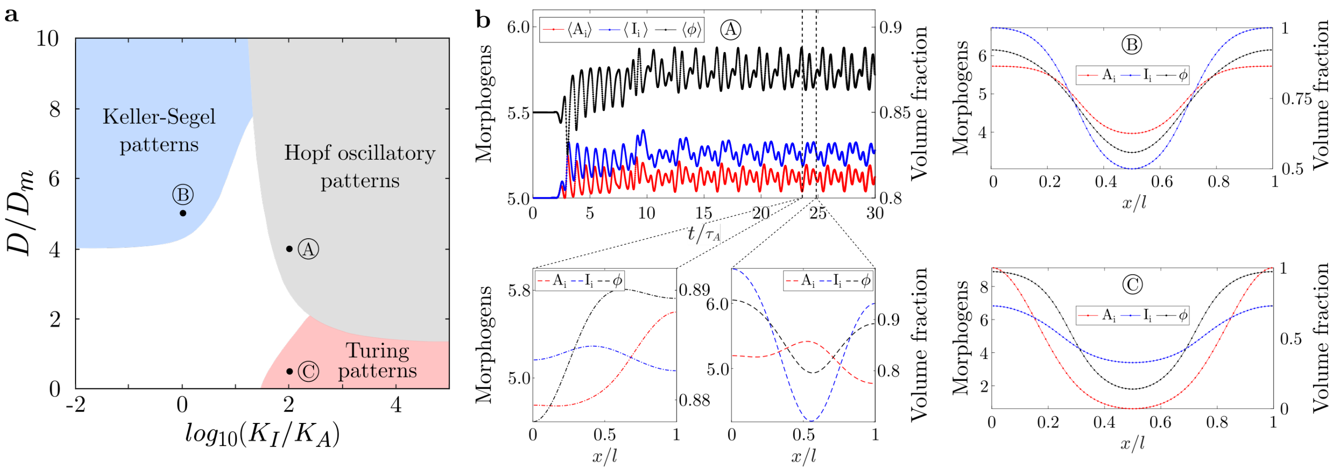


Figure S1: **Pattering instabilities of the simplified model.** (**a**) Phase diagram of (10) in the $(D_m/D, K_I/K_A)$ parameter space. In blue and red, domains corresponding to pitchfork (first and second order) bifurcations found for (10) and being respectively Keller-Segel and Turing instabilities. In grey, domain corresponding to a Hopf bifurcation and as such associated with an oscillatory instability. Model parameters are set to $\chi_A = 0.25$, $\chi_I = 1$, $\tau/(K_A\tau_A) = 1$, $\tau_I/(K_A\tau_A) = 0.2$, $K_A\tau_A\rho = 1$, $\phi^* = 0.85$ and $l_A/l \ll 1$. (**b**) 1D numerical simulations of (10) for points A, B and C, in the $(D_m/D, K_I/K_A)$ space, as shown in (a). For point A, in the Hopf oscillatory patterns region, spatially averaged morphogens concentrations and volume fraction (denoted with $\langle\rangle$) do not converge to a constant value but oscillate in time. These temporal oscillations are associated with a periodic oscillation of morphogens and volume fraction spatial profiles. Initial conditions are taken at random and $l_A/l = 0.1$.

For Fig. 4 of the main text, to illustrate the instability induced by the hydrodynamic cross-diffusion (see section "Cross-diffusion Turing instabilities" in the main text), we construct a very simple inhibitor-inhibitor scheme where:

$$\begin{aligned} f(A_i, I_i) &= -\frac{(A_i - A_i^*)}{\tau_A} - \frac{(I_i - I_i^*)}{\tau_I} + \Psi(A_i, I_i) \\ g(A_i, I_i) &= -\frac{(A_i - A_i^*)}{\tau_I} - \frac{(I_i - I_i^*)}{\tau_A} + \Psi(A_i, I_i), \end{aligned}$$

and,

$$\begin{aligned} \Psi(A_i, I_i) = -\alpha[((A_i - A_i^*)/A_i^*)^2 + ((I_i - I_i^*)/A_i^*)^2] \\ H(A_i - A_{\min})H(I_i - I_{\min}) \end{aligned}$$

is a non-linear function chosen to ensure that $A_i$ and $I_i$ remain bounded and positive ($H$ denotes the Heaviside function). Parameters used for the figure numerical simulation are $\alpha K_A \tau_A = 1$ $A_{\min}/A_i^* = I_{\min}/I_i^* = 1$ and $l_A/l = 0.1$.

Generically, $\phi_h(A_i, I_i)$ could be a rather complex function. However, for a linear stability analysis of patterning instabilities, we can restrict its functional form to a first-order linear expansion of the cell volume fraction around the reference state:

$$\phi_h(A_i, I_i) = \phi^* + \chi_A(A_i - A_i^*)/A_i^* + \chi_I(I_i - I_i^*)/I_i^* \quad (9)$$

The term $\phi^*$ accounts for mechanisms controlling cell volume and proliferation that are independent of morphogens levels, such as mechanical stress caused by active cell contractility [36], or osmotic effects due to passive water flows trough aquaporins [20, 21] and ions efflux/influx by transmembrane ion channels and pumps [6]. The $\chi_{A,I}$ terms account for the sensitivity of homeostatic cell volume fraction to intracellular morphogens concentrations and thus depend on how morphogens regulate both the volume of individual cells but also their proliferation. We also note that although this linear expansion is valid to study the onset of the instability, this does not constrain $\phi$ to be bound between 0 and 1, which is required from a physical perspective, and arises from non-linear terms. This becomes important for numerical simulations, where non-linear terms are crucial to stabilize the amplitude of the instability, so we chose for all numerical integrations:

$$\phi_h(A_i, I_i) = \frac{\phi^*\left(\tanh\left(\frac{\chi_A(A_i - A_i^*)/A_i^* + \chi_I(I_i - I_i^*)/I_i^*}{2(1-\phi^*)\phi^*}\right) + 1\right)}{(2\phi^* - 1)\tanh\left(\frac{\chi_A(A_i - A_i^*)/A_i^* + \chi_I(I_i - I_i^*)/I_i^*}{2(1-\phi^*)\phi^*}\right) + 1},$$

which is compatible with the linear approximation mentioned above.

#### 1.3 Simplified model

To achieve mathematical tractability and discuss the solution of the model in a biophysically relevent context, it is possible to reduce (6) by assuming that the import rates $\mathbb{\Gamma} \gg \mathbb{\Lambda} \gg \mathbb{J}$ are much larger than the export rates, such that $\mathbb{K} \ll 1$ is small and that the import/export dynamics is faster than morphogens turnover. From a biological standpoint (see also Section 1.7.3 for further details) $\mathbb{\Gamma}, \mathbb{\Lambda} \approx 10^{-2}\ \mathrm{s}^{-1} \gg \mathbb{J} \approx 10^{-4}\ \mathrm{s}^{-1}$, [37] and we explain in Section 1.6 why we additionally consider the limit $\mathbb{K} \ll 1$. Note however that we demonstrate in Section 1.7.1 that the Keller-Segel instability that we disclose in the single morphogen case does not depend on the simplifying double limit considered in this Section.

When $\mathbb{\Gamma} \gg \mathbb{\Lambda} \gg \mathbb{J}$, the morphognens import and export almost compensate each other such that $\underline{X}_e \simeq \mathbb{K}\underline{X}_i$. As a result the external concentrations of morphogens are small compared to the internal ones. However, exchange terms in (6) remain finite because $\mathbb{\Gamma}^{-1}(\underline{X}_e - \mathbb{K}\underline{X}_i)$ is an indeterminate product between a large quantity ($\mathbb{\Gamma}^{-1}$) and a small one ($\underline{X}_e - \mathbb{K}\underline{X}_i$). To resolve this indeterminate form, we sum up the first and second equations of (6) such that the exchange terms exactly cancel out and, as

$\underline{X}_e \simeq \mathbb{K}\underline{X}_i$, we obtain a closed system of the internal morphogens concentration:

$$\partial_t(\phi\underline{X}_i) + \nabla.\left(D_m\mathbb{K}\nabla\phi\underline{X}_i - \mathbb{K}D\nabla\underline{X}_i\right) = \underline{F}(\underline{X}_i)$$
$$\tau\partial_t\phi - \nabla.(\tau D_m\nabla\phi) = \phi_h(\underline{X}_i) - \phi.$$

For such system to be non-trivial, we assume that both $D_m\mathbb{K}$ and $\mathbb{K}D$ remain finite. This implies that the diffusion $D_m$ is large and for $l_m = \sqrt{D_m\tau}$ to remain finite as well, we also need to assume that $\tau \sim \tau_A K_A \ll 1$ is small. As a result, the cell volume fraction relaxes infinitely fast and we finally obtain the simplified model

$$\begin{aligned}\partial_t(\phi\underline{X}_i) + \nabla.\left(D_m\mathbb{K}\nabla\phi\underline{X}_i - \mathbb{K}D\nabla\underline{X}_i\right) = \underline{F}(\underline{X}_i)\\ -l_m^2\nabla^2\phi + \phi = \phi_h(\underline{X}_i),\end{aligned} \quad (10)$$

which couples a reaction-drift-diffusion equation for the morphogens levels with an elliptic equation controlling the drift as a function of the morphogen levels. Such structure is reminiscent of some previous investigations which have been performed in the completely different context of cytoskeleton self-organisation [38].

At this stage, we note that the robustness of the model with respect to two hypothesis can be demonstrated. In (6), we can add to the second equation a degradation term for the external morphogen concentrations $-\mathbb{\Gamma}_{\text{ext}}\underline{X}_e$ and we can also suppose that, in the third equation $\phi(\underline{X}_i, \underline{X}_e)$ also depends on $\underline{X}_e$. As $\underline{X}_e = \mathbb{K}\underline{X}_i$, this will not change the form of (10) but only effectively rescale the expressions of $\underline{F}$ and $\phi_h$.

Importantly, when $\chi_{A,I} = 0$ (no sensitivity of the volume fraction on the morphogen concentrations), the second equation uncouples from the first one such that $\phi \equiv \phi^*$ and (10) reduces to the classical Turing case

$$\phi^*\partial_t\underline{X}_i - \mathbb{K}D\nabla^2\underline{X}_i = \underline{F}(\underline{X}_i)$$

which is fully studied in [39] with the only difference that the global diffusion of morphogens is now rescaled by the transmembrane transport equilibrium constants (see section "Orders of magnitude on morphogen transport" in the main text).

#### 1.4 Stability analysis of the simplified model

We are now interested in determining if we can obtain a stationary spatial pattern for morphogens concentrations in perturbing the system (10) in the vicinity of its homogeneous equilibrium state. To do so, we write:

$$\underline{X}_i = \underline{X}_i^* + \epsilon\delta\underline{X}_i \text{ and } \phi = \phi^* + \epsilon\delta\phi$$

where $\epsilon \ll 1$, and we obtain at the first order in $\epsilon$:

$$\underline{X}_i^*\partial_t\delta\phi + \phi^*\partial_t\delta\underline{X}_i + D_m\mathbb{K}\underline{X}_i^*\nabla^2\delta\phi - D\mathbb{K}\nabla^2\delta\underline{X}_i = \mathbb{J}\delta\underline{X}_i$$
$$-l_m^2\nabla^2\delta\phi + \delta\phi = \nabla_{\underline{X}_i}\phi_h|_{\underline{X}_i^*}.\delta\underline{X}_i.$$

We now introduce the sets of eigenvalues $\{\lambda_k\}_{k\geq 1}$ and eigenvectors $\{U_k(x)\}_{k\geq 1}$ of the minus Laplace operator defined by the problem:

$$\begin{cases} -\nabla^2 U_k = \lambda_k U_k \\ \nabla U_k.n|_{\partial\Omega} = 0. \end{cases} \quad (11)$$

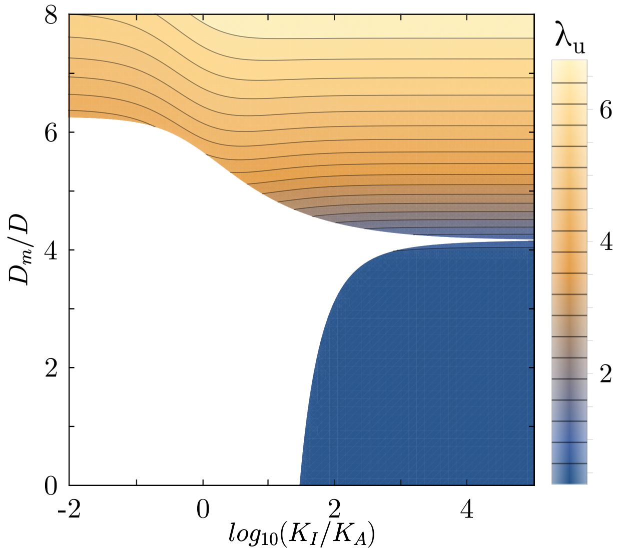


Figure S2: **Estimation of the pattern wavelength using linear stability.** Contour plot of the most unstable wavelength obtained using the dispersion relation (10) in the $(K_I/K_A, D_m/D)$ parameter space shown on Fig. 2 (a) of the main text. parameters are set to $\tau/(K_A\tau_A) = 0.01$, $\chi_A = 0.25$, $\chi_I = 0$, $\tau_I/(K_A\tau_A) = 0.2$, $K_A\tau_A\rho = 1$, $\phi^* = 0.85$ and large tissue size $(l_A/l \ll 1)$.

By classical theorems [39], $\{\lambda_k\}$ is a countable set of positive and increasing values, and $\{U_k\}$ forms a basis on which we can project the solutions of the linearised system :

$$\delta\underline{X}_i(\vec{r}, t) = \sum_{k=1}^{\infty} \underline{X}_i^k U_k(\vec{r})\exp(\omega_k t),$$

where $\omega_k$ is the growth rate associated to the $k^{\text{th}}$ growth mode. For example, when $\Omega$ is a one dimensional segment of size $l$ we have $\lambda_k = (\pi k/l)^2$ and $U_k(x) = \cos(\pi kx/l)$. From this ansatz, we obtain that:

$$\left[\omega_k\left(\frac{\mathbb{C}}{1+\lambda_k l_m^2} + \phi^*\mathbb{I}\right) + \lambda_k\mathbb{K}\left(D\mathbb{I} - \frac{D_m\mathbb{C}}{1+\lambda_k l_m^2}\right) - \mathbb{J}\right]\underline{X}_i^k = 0$$

where,

$$\mathbb{C} = \begin{pmatrix} \chi_A & \chi_I r^{-1} \\ \chi_A r & \chi_I \end{pmatrix} \text{ with } r = \frac{A_i^*}{I_i^*}.$$

Thus, unstable spatial modes are those for which a $\omega_k$ with a positive real part solves the dispersion relation:

$$\det\left[\omega_k\left(\frac{\mathbb{C}}{1+\lambda_k l_m^2} + \phi^*\mathbb{I}\right) + \lambda_k\mathbb{K}\left(D\mathbb{I} - \frac{D_m\mathbb{C}}{1+\lambda_k l_m^2}\right) - \mathbb{J}\right] = 0. \quad (12)$$

To be in the same case as in Turing's model, we impose the constraint that the homogeneous solution in the absence of spatial terms ($\lambda_k = 0$) is stable. Thus the two solutions $\omega^0$ of

$$\det\left[\omega^0\left(\mathbb{C} + \phi^*\mathbb{I}\right) - \mathbb{J}\right] = 0,$$

have negative real parts. Such condition is satisfied as soon as

$$\phi_* + \text{tr }\mathbb{C} > 0, \ \det\mathbb{J} > 0 \text{ and } \phi_*\text{tr }\mathbb{J} + \det\mathbb{J}\text{tr }(\mathbb{J}^{-1}\mathbb{C}) < 0. \quad (13)$$

Otherwise, the homogeneous solution can lose stability even without spatial terms and the steady state would then oscillate in time. These conditions generalize the classical ones [39], det $\mathbb{J} > 0$ and tr $\mathbb{J} < 0$ obtained for $\chi_{A,I} = 0$.

In the main text (See Fig 2(a)), we study the dispersion relation (12) in the case where $\mathbb{J}$ is computed based on the classical activator-inhibitor scheme (8) and $\chi_I = 0$ since conditions (13) are automatically satisfied in this case. With such choice, the instability of the homogeneous state is a pitchfork bifurcation, meaning that at the onset of instability, the imaginary part of the unstable(s) mode(s) $\omega_k$ is zero. Among the unstable $\lambda_k$ (i.e. those associated with a positive $\omega_k$), we can select the most unstable one (i.e. the one associated with the largest value of $\omega_k$) in the dispersion relation (12). This wavelength $\lambda_u$ is the one which grows with the largest rate and is therefore likely to be the observed one in a numerical simulation of (10). However, this result should not be considered as rigorous given that there may be other non-linear effects selecting the wavelength which escape this analysis.

To complement Fig 2(a) of the main text, we show on Fig. S2 a contour plot of the values of $\lambda_u$ in the ($K_I/K_A$, $D_m/D$) phase space. In the Turing limit (captured by $D_m/D \ll 1$ on Fig. 2(a) of the main text), $\lambda_u$ depends very weakly on the ratio $K_I/K_A$ given that this ratio has to be large for the instability to occur in the first place. Therefore $\lambda_u$ is mostly controlled by the turnover rates (fixed in Fig. S2) in this case. In the Keller-Segel limit (captured by $K_I/K_A \ll 1$ on Fig 2(a) of the main text), the most unstable wavelength depends on both the turnover rates (fixed) and the ratio $D_m/D$ which explains its variation along this axis. Note that the number of patterns shown on direct numerical simulations shown on Fig 2(b) of the main text satisfy this linear prediction of the total number of patterns because they are close to the onset of instability.

To complement this analysis, we also show that considering non-vanishing $\chi_I$, but remaining within the admissible bounds of (13), can lead to the existence of a third phase representing a Hopf bifurcation where the solution becomes oscillatory. Although interesting, this goes beyond the scope of our analysis, which concentrates on steady-state spatial patterns, and live-imaging would be needed to verify whether complex time-oscillations can occur during patterning in developmental settings. We illustrate however this dynamic behavior in Fig. S1, in which we also show direct numerical simulations of (10) to justify the claim. We did not find other type of instabilities than the Turing, Keller-Segel and oscillatory ones regardless of the sign of $\chi_A$ and $\chi_I$ in the case of a classical activator-inhibitor scheme.

#### 1.5 Cross-diffusion Turing patterns

An other case of special interest is when cell volume fraction sensitivities to morphogen concentration are negative ($\chi_{A,I} < 0$), eliminating the possibility of up-hill morphogen diffusion at the origin of the Keller-Segel instability in (12).

To simply explain the physical nature of the instability developing then, we first set $\tau = 0$ (leading to $l_m = 0$) such that the tissue volume fraction always locally remains at its homeostatic value. We also assume that the sensitivities of the volume fraction are small $\chi_{A,I} \ll 1$, but that the effective fickian and hydrodynamic diffusivities $D\mathbb{K} \sim D_m\mathbb{K}\mathbb{C}$ remain finite and of the same order of magnitude. Here the aim is not to quantatively justify these limits from a biological standpoint, given the lack of quantitative measurements of $\chi_{A,I}$, but our goal is to give a qualitative explanation of the instability in a mathematically tractable setting (although the core mechanism we describe does not depend on this assumption).

In this limit, the conditions for linear stability of the homogeneous system (13) reduces to the classical ones while the dispersion relation (12) reads:

$$\det\left[\omega_k \phi^* \mathbb{I} + \lambda_k \mathbb{K}\left(D\mathbb{I} - D_m \mathbb{C}\right) - \mathbb{J}\right] = 0. \tag{14}$$

Interestingly, this relation of dispersion has been studied for monophasic reaction-diffusion systems with *ad hoc* cross-diffusion terms [40]. We also refer to [41] for further exposure of the potential physical origin of cross-diffusion terms in reaction-diffusion system. As shown in [40], such cross diffusion terms result in a dramatic broadening of the patterns phase space. In particular, any two-morphogen reaction scheme has the potential to generate spatial patterns and not just the classical activator-inhibitor schemes.

#### 1.6 Generalized Turing instability case

We now get back to the general case of system (6) but assume that $\chi_{A,I} = 0$ such that cell volume fraction is not sensitive to morphogen concentration. In this case, the last equation uncouples and relaxes to $\phi \equiv \phi^*$ such that (6) reduces to,

$$\begin{aligned} \phi^* \partial_t \underline{X}_i &= \underline{F}(\underline{X}_i) + \mathbb{\Gamma}\underline{X}_e - \mathbb{\Lambda}\underline{X}_i \\ (1-\phi^*)\partial_t \underline{X}_e - D\nabla^2 \underline{X}_e &= -\mathbb{\Gamma}\underline{X}_e + \mathbb{\Lambda}\underline{X}_i \end{aligned} \tag{15}$$

The linear stability of the homogeneous solution $\underline{X}_i \equiv \underline{X}_i^*$, $\underline{X}_e \equiv \mathbb{K}\underline{X}_i^*$ of (15) reduces to the dispersion relation

$$\det\left[\left(\omega_k(1-\phi^*) + D\lambda_k\right)\left(\omega\phi^*\mathbb{I} - \mathbb{J} + \mathbb{\Lambda}\right) + (\omega\phi\mathbb{I} - \mathbb{J})\mathbb{\Gamma}\right]. \tag{16}$$

In the main text, we consider the double limit:

$$\mathbb{\Gamma} \gg \mathbb{\Lambda} \gg \mathbb{J},$$

in which case, the above relation reduces to the classical [39] Turing case,

$$\det\left[\omega\phi\mathbb{I} + \lambda_k D\mathbb{K} - \mathbb{J}\right],$$

and the analogue of (15) reads:

$$\phi^* \partial_t \underline{X}_i - D\mathbb{K}\nabla^2 \underline{X}_i = \underline{F}(\underline{X}_i).$$

While it is clear from a biological standpoint that $\mathbb{\Gamma}, \mathbb{\Lambda} \approx 10^{-2}\ \mathrm{s}^{-1} \gg \mathbb{J} \approx 10^{-4}\ \mathrm{s}^{-1}$, [37], it is not clear that $\mathbb{\Gamma} \gg \mathbb{\Lambda}$. However, we note that the Turing instability typically holds in this limiting case. As a matter of fact,

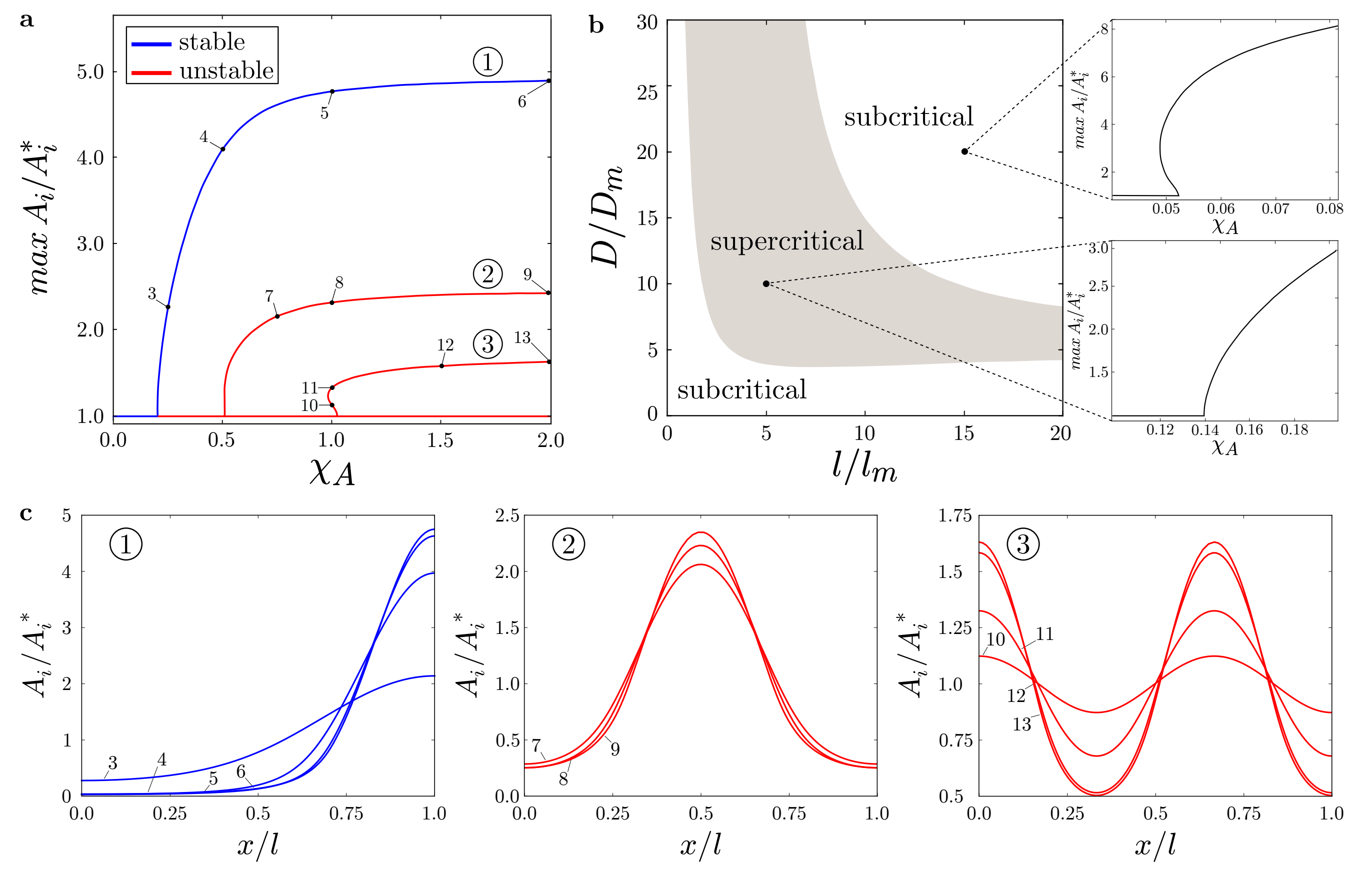


Figure S3: **Bifurcation diagram of the solution** (22) **of the simplified model with one morphogen.** (**a**) First branches bifurcating from the homogeneous solution at the onset of instability. Black lines are stable, while red lines are unstable. Stability is obtained numerically in integrating the time dependent problem (21), starting from the considered steady state solution with a small perturbation. Parameters are $\phi^* = 0.85$, $D_m/D = 10$ and $l/l_m = 3.1$. (**b**) Nature of the fundamental (i.e. first) bifurcation point. Parameter $\phi^* = 0.85$. (**c**)Profiles of activator concentrations along the branches shown in (**a**)(*cf.* corresponding labels). Half of a wavelength is added each time the index of the branch is increased by one. Increase in $\chi_A$ results in sharper profiles. Only profiles along one branch of the pitchfork bifurcation are shown. The other branch, superimposed on the one shown, corresponds to symmetric profiles in respect with the centre of the space domain.

the opposite limit where $\mathbb{A} \gg \mathbb{\Gamma}$ does not lead to patterns. Indeed, in the case

$$\mathbb{A} \gg \mathbb{\Gamma} \gg \mathbb{J},$$

(16) reduces to:

$$\det\left[\omega(1-\phi^*)\mathbb{I} + \lambda_k D\mathbb{I} - \mathbb{J}\mathbb{K}^{-1}\right],$$

which cannot lead to a patterning instability in general as it corresponds to a case where the diffusion coefficients for the activator and the inhibitor are the same.

#### 1.7 Single morphogen case

We now consider the general case where a single morphogen exists which we chose by convention to be $A$. In that case, (6) simplifies to:

$$\begin{gathered}\partial_t(\phi A_i) = f(A_i) + \gamma_A A_e - \lambda_A A_i \\ \partial_t\left((1-\phi)A_e\right) + \nabla.\left(A_e D_m \nabla\phi - D\nabla A_e\right) = -\gamma_A A_e + \lambda_A A_i \\ \partial_t\phi - \nabla.(D_m\nabla\phi) = \frac{\phi_h(A_i)-\phi}{\tau}.\end{gathered} \tag{17}$$

where $f(A_i) = (A_i^* - A_i)/\tau_A$ is the linearized form of $f$.

##### 1.7.1 Robustness with respect to the main text limit

In the main text, system (17) is studied in the limit where $\gamma_A \gg \lambda_A \gg 1/\tau_A$ and $\tau/\tau_A \sim K_A \ll 1$ and thus reduces to:

$$\begin{gathered}\partial_t(\phi A_i) + K_A\nabla.\left(A_i D_m\nabla\phi - D\nabla A_i\right) = f(A_i) \\ -l_m^2\nabla^2\phi + \phi = \phi_h(A_i).\end{gathered} \tag{18}$$

However, an explicit criterion for the linear stability of (17) can be obtained where instead of such complex limit, we simply postulate (only for sake of simplicity) that $\phi^* \simeq 1$. While $\gamma_A, \lambda_A \gg 1/\tau_A$ is verified in biological situations, this is particularly helpful to demonstrate that the instability we find in the single morphogen case does not rely on the assumption $\gamma_A \gg \lambda_A$. The condition for an instability to occur in (17) (when the characteristic domain size $l \gg \sqrt{D\tau_A}$) reads,

$$\sqrt{\frac{D}{D_m}\left(1 + \frac{1}{K_A\gamma_A\tau_A}\right)} + \sqrt{\frac{\tau}{K_A\tau_A}} < \sqrt{\chi_A}, \tag{19}$$

which can be further reduced to the condition formulated in the main paper:

$$\sqrt{\frac{D}{D_m}} + \sqrt{\frac{\tau}{K_A\tau_A}} < \sqrt{\chi_A}, \tag{20}$$

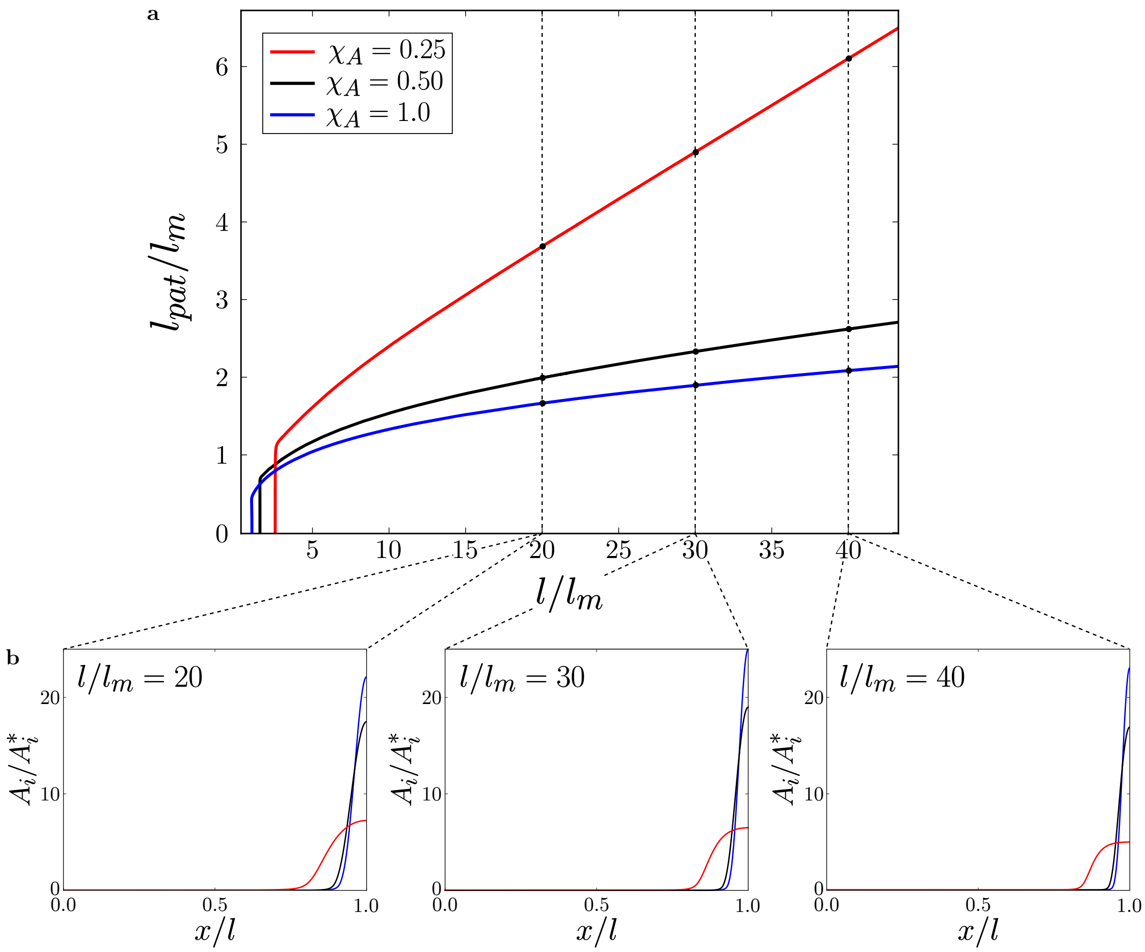


Figure S4: **Scaling of pattern size with domain size, for the simplified model with one morphogen.** (**a**) Size of the pattern as a function of tissue domain size. After the onset of instability, the stable pattern size increases linearly with the domain size for various values of $\chi_A$. Parameters are $D_m/D = 10$ and $\phi^* = 0.85$. (**b**) Morphogen (activator) concentration profiles along selected point of the branches shown in panel (**a**).

in the biologically meaningful case (see Section 1.7.3) where $K_A\gamma_A\tau_A \gg 1$. To further demonstrate this claim we present on Fig. S4 direct numerical simulations displaying steady state patterns for large and small values of $K_A$.

##### 1.7.2 Scaling properties of Keller-Segel patterns

In this section, we consider only the special case where $\tau_A \to \infty$ and $K_A \to 0$ while $K_A D_m$, $K_A D$ and $\tau D_m$ remain finite. This limit corresponds to a case where the morphogen turnover rate is very slow but where the effective fickian and hydrodynamic diffusivities remain nonetheless finite. In such case, (17) reduces to a classical Keller-Segel system [42]:

$$\begin{aligned} \partial_t(\phi A_i) + K_A \nabla.\left(A_i D_m \nabla\phi - D\nabla A_i\right) = 0 \\ -l_m^2 \nabla^2\phi + \phi = \phi_h(A_i), \end{aligned} \tag{21}$$

but with a slightly different dynamics ($\phi A_i$ instead of $A_i$ in the time derivative) and a non-linear dependence of $\phi$ on $A_i$ through $\phi_h$. A very similar equation was derived and studied in the context of active gels [38, 43].

Integrating the first equation of (21), steady states of (21) are given by

$$-l_m^2\nabla^2\phi + \phi = \phi_h\left(A_i^* \frac{\mathrm{e}^{\frac{D_m}{D}\phi(\vec{r})}}{\left\langle \mathrm{e}^{\frac{D_m}{D}\phi}\right\rangle}\right), \tag{22}$$

where $\langle h\rangle = |\Omega|^{-1}\int_\Omega h(\vec{r})d\vec{r}$ denotes the spatial average over the tissue domain $\Omega$. For simplicity, we now consider (22) on the finite segment $[0, l]$ for which $\lambda_k = (\pi k/l)^2$, and investigate the possible bifurcations arising from the homogeneous state when the volume fraction sensitive parameter $\chi_A$ increases. Using the continuation software AUTO [44], we numerically compute the first three branches of such bifurcation diagram on Fig. S3 (a). It features an infinite number of critical points $\chi_A^k = D/D_m(1 + l_m^2\lambda_k)$ from where inhomogeneous solutions bifurcate, see Fig. S3 (c).

Similar to [38, 43], we numerically show that starting from an arbitrary distribution of $A_i$ and if an instability exists, the system (21) will always coarsen to a single "two-zones" pattern corresponding to the fundamental mode of the instability $\lambda_1$. The reason for this is that morphogen concentration creates a non-local convergent (since $\chi_A$ positive) flow recruiting even more morphogens over the hydrodynamic length $l_m$ and further increasing the flow. This positive feedback loop would lead to infinite localization of the morphogen concentration if morphogen diffusion was not preventing such a blow up by penalizing the creation of sharp gradients in concentration. If $\chi_A$ is small, the diffusion dominates and the concentration of morphogen is homogeneous but when $\chi_A$ reaches the critical threshold $\chi_A^1$, the morphogen self-organize into a fundamental pattern where active advective transport balances diffusion. Th is is known as the Keller-Segel mechanism. Because the positive feedback loop is non-local, the morphogens concentration always coarse-grains into the fundamental mode. Interestingly, we found using a classical normal mode analysis, and confirmed numerically, that the nature of this first bifurcation branch is of pitchfork type but can be supercritical (second order phase transition) or subcritical (first order phase transition) *cf.* Fig. S3 (b). In practice, this means that two steady states corresponding to a non-patterned and a patterned state may co-exist for a range of parameters in the subcritical case. The transition from one state to the other could be then controlled by additional regulatory pathways.

Next, we define the fundamental pattern size as $l_{\text{pat}} = \int_0^l H(A_i(x) - A_i^*)dx$, where the Heaviside function selects the part of the morphogen profile larger than its average value. Therefore $l_{\text{pat}} = 0$ for the homogeneous solution and becomes non-zero when a pattern forms. We show on Fig. S4 that this pattern scales as a function of the domain size for various values of $\chi_A$. These results demonstrate that when an instability exists, a single "two zones" pattern always forms and its size scales with an increase in tissue domain size. This scaling property is at odds with what is observed for a classical Turing instability, where additional patterns form as the domain size increases. Thus, in this limit, our model constitutes a suitable theory to explain the scaling properties of various "two-zones" developmental patterns such as the dorso-ventral pattern of *Xenopus* [45], the antero-posterior pattern of planarian [46] or the left-right pattern of zebrafish [47].

##### 1.7.3 Biological relevance of the Keller-Segel instability

Here, we briefly review the orders of magnitude from the main text, to justify in particular that the Keller-Segel-Turing phase diagram proposed in Fig 2 of the main text is relevant given the reported orders of magnitude for the model parameters. We concentrate here in particular on the Keller-Segel instability and (19). The hypothesis of $K_A\gamma_A\tau_A \gg 1$ is rather straightforwardly relevant, as typical values measured in the literature are $\tau_A \approx 10^4-10^5$ s [47], while $\gamma_A \approx 10^{-1}-10^{-2}s^{-1}$ [37]. Therefore, even for small values of $K_A$, the approximation is expected to hold (as values at which it would break, $K_A < 10^{-3}$ would be when almost no morphogen is present extracellularly anymore). This shows that the instability condition derived in the main paper for the Keller-Segel instability is in fact robust with respect to the assumption that $K_A \ll 1$ (contrary to the Turing insta bility with re-normalized coefficients which require $K_A \ll 1$).

Thus, one can adopt the simplified instability criteria (20), where $\sqrt{\frac{D}{D_m}}$, $\sqrt{\frac{\tau}{K_A\tau_A}}$ and $\sqrt{\chi_A}$ must be compared. Because $\chi_A$ is the least understood of these effects, we start by concentrating on the first two. Orders of magnitude from the main text allow us to estimate $\tau/(K_A\tau_A) \approx 0.01 - 0.1$. Moreover, $D_m$ can adopt a wide range of values given the wide range of length scales of extracellular gaps in different tissues, but its value is consistently larger to much larger than $D$, with $D/D_m \approx 10^{-4}-1$, which tends to say that that $\tau/(K_A\tau_A)$ is the main factor resisting the Keller-Segel instability in the limit of large hydrodynamic diffusion $D_m$ (i.e. when extracellular pore sizes are 0.1 micron-sized). In this case, volume fraction sensitivities $\chi_A$ must be larger than $0.01-0.1$ for the Keller-Segel instability to develop, which, although relative ly little data exists on variations of $\phi$ in developing tissues, is a rather lax criterion, as it represents typical variations of $\phi$ of only $1-10\%$ with morphogen concentration. One should note that for very compact tissues, where values of $\sqrt{D/D_m}$ could be close to unity, the instability could require unrealistically large values of $\chi_A$ and would be screened.

#### 1.8 Linking FRAP and FCS measurements of the diffusion coefficient

Fluorescence recovery after photobleaching (FRAP) is a method which has been longed used to experimentally determine diffusion coefficients [48]. More particularly, it has been employed recently in several *in vivo* systems such as the early Zebrafish [47] and Xenopus [45] embryos or the Drosophila wing disk [49] in order to measure tissue-wide effective diffusion coefficients of various morphogens. Nevertheless, establishing a correspondence between these FRAP measured diffusion coefficients and the effective diffusion coefficient defined in our modeling framework requires some care.

Indeed, in such FRAP experiments, the endogeneous molecule of interest is usually replaced by a genetically engineered copy which has been fused to a fluorescent protein such as the GFP. Thus, if one wants to study simultaneously two interacting morphogens, one needs to genetically engineer two constructs with two different fluorescent protein tags. In practice this is rarely the case and the diffusive properties of each morphogen are studied idependently [45,47]. Moreover, such an approach does not directly discriminate the internal and external fractions of a given morphogen. Therefore, the intensities of fluorescence are proportional to the total concentration of a given morphogen : $A_t = \phi A_i + (1-\phi)A_e$ for $A$ and $I_t = \phi I_i + (1-\phi)I_e$ for $I$. Combining the conservation equations (2) and (3), we obtain:

$$\begin{aligned} \partial_t A_t + \nabla.(A_e D_m \nabla\phi - D\nabla A_e) &= f(A_i, I_i) \\ \partial_t I_t + \nabla.(I_e D_m \nabla\phi - D\nabla I_e) &= g(A_i, I_i) \\ \partial_t \phi - \nabla.(D_m \nabla\phi) &= \frac{\phi_h(A_i,I_i)-\phi}{\tau}. \end{aligned} \qquad (23)$$

For a small enough bleached region, the contribution of morphogen turnover rates can be neglected and the fluorescence recovery is mostly driven by transport, as shown in the case of the Zebrafish embryo [47]. For the sake of simplicity, we shall thus proceed under this assumption, although some other *in vivo* systems may require to account for morphogen turnover rates [50].

The classical Turing case corresponds to a fixed volume fraction $\phi = \phi^*$ implying $v_e = 0$. As equations for $A$ and $I$ uncouple, we can focus on a single morphogen only (say

$A$) and the fluorescence $A_t$ satisfies:

$$\begin{array}{c}\partial_t A_t - \nabla.(D\nabla A_e) = 0\\ \phi^*\partial_t A_i = \gamma_A A_e - \lambda_A A_i\\ A_t = \phi^* A_i + (1-\phi^*)A_e.\end{array} \quad (24)$$

From (24), we can deduce an dispersion relation

$$(1-\phi^*)\omega^2 + \left(\gamma_A + \lambda_A\frac{1-\phi^*}{\phi^*} + D\lambda_k\right)\omega + \frac{\lambda_A D\lambda_k}{\phi^*} = 0$$

where $\lambda_k$ are the eigenvalues of the Laplace operator over the bleached region of typical size $L_b$ and $\omega$ the rates associated with fluorescence recovery. Non dimensionalizing times with $\gamma_A^{-1}$ and introducing the non-dimensional quantity $\Delta = D/(\gamma_A L_b^2)$ we can rewrite the above equation as

$$(1-\phi^*)\bar{\omega}^2 + \left(1 + K_A\frac{1-\phi^*}{\phi^*} + \rho_k\Delta\right)\bar{\omega} + \frac{K_A\rho_k\Delta}{\phi^*} = 0 \quad (25)$$

where $\bar{\omega}$ are non-dimensional rates and $\rho_k = L_b^2\lambda_k$ is a positive non-dimensional geometric factor which increases quadratically with the order of the eigenvalue considered. For instance if the bleached region is spherical $\rho_k$, is the discrete set of zeros of the Bessel functions of the first kind [48]. If the bleached region is $\sim 100\ \mu$m [47], we have that $L_b^2\gamma_A \sim 10^{-9}\text{m}^2\text{s}^{-1}$ suggesting that $\Delta \ll 1$ in (25). Then at first order we have

$$\bar{\omega}_1 = -\frac{K_A(1-\phi^*)+\phi^*}{\phi^*(1-\phi^*)} \text{ and } \bar{\omega}_2 = -\frac{K_A\rho_k\Delta}{K_A(1-\phi^*)+\phi^*},$$

where $\bar{\omega}_1 \gg \bar{\omega}_2$ is an homogeneous clearance rate of fluorescence that is important only at very short timescales while $\bar{\omega}_2$ corresponds to the measured diffusion driven recovery with effective diffusion coefficient,

$$D_{\text{FRAP}} = \frac{K_A D}{K_A(1-\phi^*)+\phi^*}.$$

To relate the effective diffusion coefficient $D_{\text{FRAP}}$ to the local diffusion coefficient $D_{\text{Fick}}$ measured by Fluorescence Correlation Spectroscopy (FCS), one should recall that $D$ is already a homogenized over a RVE and as such, depends on the tissue packing fraction. A classical way to relate $D$ and $\phi^*$ is through the Maxwell formula [51]:

$$D \simeq D_{\text{Fick}}\frac{1-\phi^*}{1+\phi^*/2}.$$

Maxwell's result formally only holds in the dilute approximation where $\phi^*$ is close to zero but in fact corresponds to the Hashin-Shtrikman [52] upper-bound which is known to hold outside the dilute limit:

$$D \leq D_{\text{Fick}}\frac{1-\phi^*}{1+\phi^*/2}.$$

Such bound can be computed irrespective of the microscopic details of tissue geometry meaning that the actual effect of tortuosity is in reality even more pronounced. One can then directly compare the diffusion coefficient measured by FRAP and the one measured by FCS. For instance, in our first case where $\Delta \ll 1$, we obtain:

$$\frac{D_{\text{FRAP}}}{D_{\text{Fick}}} \leq \frac{K_A(1-\phi^*)}{(\phi^*+K_A(1-\phi^*))(1+\phi^*/2)}.$$

Interestingly, Lefty diffusion coefficient in the zebrafish embryo varies only by a factor 2 between FCS and FRAP measurements [47], $D_{\text{FRAP}}/D_{\text{Fick}} \simeq 0.5$. This can hold in the limit of $K_{\text{Lefty}} \gg 1$, since in this case we obtain $D_{\text{FRAP}} \leq \frac{D_{\text{Fick}}}{1+\phi^*/2}$, which is consistent with the statement of Muller et al. in [47] that there is "an approximately two- to three-fold decrease in diffusivity in the presence of cells compared to unhindered diffusion in aqueous solution". Note that this last bound ($K_{\text{Lefty}} \gg 1$ limit) also corresponds to the one expected for a passive extracellular molecule which does not chemically interact with cells such as an injected dye or a secreted recombinant fluorescent protein (see [50] for experiments of this kind in early Zebrafish embryos). However, if $K_{\text{Lefty}} \lesssim 1$ (concentrations outside of the same order of magnitude or smaller than concentration inside or immobilized), given the large cell volume fraction inferred in this system ($\phi^* \approx 0.8-0.9$) [47], we expect the decrease in effective diffusion due to tortuosity to become much larger: $D_{\text{FRAP}}/D_{\text{Fick}} \leq 0.1$. This argues for the need for precise measurements of $K_{\text{Lefty}}$ concomitant to $\phi$. Another possibility is to consider the possibility of active contributions to morphogen transport ($v_e \neq 0$ in (23)) since hydrodynamical diffusion would then, to leading order, rescale passive diffusion in an arbitrary manner as we show below.

If $\phi$ is no longer assumed to be homogeneous, variations of $\phi$ may contribute to the fluorescence recovery of the bleached region and one needs to fit the intensities $A_t$ and $I_t$ in (23) (with $f = g = 0$). A quantitative re-interpretation of the measurements of [50] in this context requires the spatial knowledge of $\phi$ and is thus left to future work. As a proof of principle, a simple case can be investigated where $\chi_I = 0$ and $\tau \to 0$. In this situation (24) becomes

$$\begin{array}{c}\partial_t A_t + \nabla.(A_e D_m\nabla\phi - D\nabla A_e) = 0\\ \partial_t(\phi A_i) = \gamma_A A_e - \lambda_A A_i\\ A_t = \phi A_i + (1-\phi)A_e\\ \phi - \phi^* = \chi_A\frac{A_i - A_i^*}{A_i^*}\end{array}$$

which at linear order reads:

$$\begin{array}{c}\partial_t A_t + \nabla.(K_A\chi_A D_m\nabla A_i - D\nabla A_e) = 0\\ (\chi_A+\phi^*)\partial_t A_i = \gamma_A(A_e - K_A A_i)\\ A_t = (\phi^* + \chi(1-K_A))A_i + (1-\phi^*)A_e.\end{array} \quad (26)$$

The generalized dispersion relation then reads:

$$\begin{array}{c}(1-\phi^*)(\chi_A+\phi^*)\omega^2 + [(\chi_A+\phi^*)(\gamma_A + D\lambda_k)\\ +\lambda_A(1-\phi^*-\chi_A)]\omega + \lambda_A\lambda_k(D - D_m\chi_A) = 0.\end{array}$$

Next, assuming $\chi_A \ll 1$ while $\chi_A D_m \sim D$ are of the same order of magnitude, we can generalize the previous result obtained for $\chi_A = 0$ to

$$D_{\text{FRAP}} = \frac{K_A(D - \chi_A D_m)}{K_A(1-\phi^*)+\phi^*}.$$

This shows that the active hydrodynamic transport could contribute to rationalize the diffusion coefficients measured by FRAP. In particular, the inequality between $D_{\text{FRAP}}$ and $D_{\text{Fick}}$ becomes

$$D_{\text{FRAP}} \leq \frac{K_A(D_{\text{Fick}} - \chi_A K l_i^2/\eta)(1-\phi^*)}{(K_A(1-\phi^*)+\phi^*)(1+\phi^*/2)},$$

which could explain, among other possibilities of active morphogens transport [53], why in some cases even with $\phi^* \simeq 0.85$ and $K_A \lesssim 1$, $D_{\text{FRAP}}$ is only slightly smaller than $D_{\text{Fick}}$.

#### 1.9 Generalisation of the model in presence of tissue rearrangements

In model (6), we have systematically assumed that changes of the tissue topology are much slower than the fluid circulation ($v_s \simeq 0$) based on the estimates presented in Section 1.1.3. However, these estimates may substantially vary depending on the biological system and cellular flows have been shown to take place at timescales comparable to fluid flows for various morphogentic processes [54]. We therefore present below an extension of the theory accounting for such flows.

A first consequence of a non-negligible $v_s$ was already discussed in Section 1.1.6 as (3) now contains a transport term

$$\begin{aligned}\partial_t(\phi A_i) + \nabla(\phi A_i v_s) = f(A_i, I_i) + \gamma_A A_e - \lambda_A A_i \\ \partial_t(\phi I_i) + \nabla(\phi I_i v_s) = g(A_i, I_i) + \gamma_I I_e - \lambda_I I_i.\end{aligned} \quad (27)$$

The next step is to obtain the expression of $v_e$ as it enters in (1) and (2). For this, we have to generalize the mechanical framework presented in Section 1.1.7 to obtain the analogue of (5) when the tissue can rearrange. We can do so by introducing a new variable $\mathbb{E}_s$ accounting for the non elastic strain present in the tissue due to the cells rearagements [55]. Thus, this strain is connected to the cell flow through the kinetic relation:

$$\frac{d\mathbb{E}_s}{dt} = \partial_t \mathbb{E}_s + v_s.\nabla\mathbb{E}_s = \nabla v_s. \quad (28)$$

In this case, the constitutive behavior (4) becomes

$$\begin{aligned}\sigma = 2G(\mathbb{E}-\mathbb{E}_s) + \left(K - \tfrac{2G}{3}\right)\text{tr}(\mathbb{E}-\mathbb{E}_s)\mathbb{I} - \alpha p \mathbb{I} \\ p = -\tfrac{K_u-K}{\alpha}\text{tr}(\mathbb{E}-\mathbb{E}_s) - \tfrac{K_u-K}{\alpha^2}\delta\phi - (1-\phi^*)\tfrac{K_u-K}{\alpha^2}\text{tr}(\mathbb{E}_s).\end{aligned} \quad (29)$$

If we continue to assume that the interstitial fluid flow in the tissue is described by Darcy's law and that the physiologically relevant limit $K_u \gg K \gg G$ and $\alpha \simeq 1$ still holds, we obtain using force balance that:

$$(1-\phi)(v_e - v_s) = -\frac{\kappa}{\eta}\nabla p = \frac{K\kappa}{\eta}\left(\nabla\phi + (1-\phi^*)\nabla\text{tr}(\mathbb{E}_s)\right).$$

Eliminating $v_e$ and denoting $\epsilon_s = \text{tr}(\mathbb{E}_s)$ we can rewrite the full model as

$$\begin{aligned}\partial_t(\phi \underline{X}_i) + \nabla.(\phi \underline{X}_i v_s) = \underline{F}(\underline{X}_i) + \mathbb{\Gamma}\underline{X}_e - \mathbb{\Lambda}\underline{X}_i \\ \partial_t((1-\phi)\underline{X}_e) + \\ \nabla.\left((1-\phi)\underline{X}_e v_s + D_m(\nabla\phi + (1-\phi^*)\nabla\epsilon_s)\underline{X}_e - D\nabla\underline{X}_e\right) \\ = -\mathbb{\Gamma}\underline{X}_e + \mathbb{\Lambda}\underline{X}_i \\ \partial_t\phi - \nabla.\left((1-\phi)v_s + D_m(\nabla\phi + (1-\phi^*)\nabla\epsilon_s)\right) = \tfrac{\phi_h(\underline{X}_i)-\phi}{\tau} \\ \partial_t\epsilon_s + v_s.\nabla\epsilon_s = \nabla.v_s.\end{aligned} \quad (30)$$

Such model generalizes (6) in the presence of a given cellular flow $v_s$. Large scale cellular flows are ubiquitous during developmental morphogenesis [56] but their genesis is far from being understood and a general assessment of their influence on our system is not presently within reach. Thus, for the sake of clarity and to extract the key biophysical ingredients generating the instabilities described in the main text, we have decided to work in the limit where $v_s = 0$.

However, we first note that if the cell flow is not associated with volumetric changes in the tissue (i.e. $\nabla.v_s = 0$) as it has been verified for example during the ventral furrow formation of Drosophilia [57] or in limb bud morphogenesis [58], then $\epsilon_s = 0$. In this case, the homogeneous solution (7) is still a solution of (30) which can be rewritten,

$$\begin{aligned}\tfrac{d\phi\underline{X}_i}{dt} = \underline{F}(\underline{X}_i) + \mathbb{\Gamma}\underline{X}_e - \mathbb{\Lambda}\underline{X}_i \\ \tfrac{d(1-\phi)\underline{X}_e}{dt} + \nabla.\left(D_m\nabla\phi\underline{X}_e - D\nabla\underline{X}_e\right) = -\mathbb{\Gamma}\underline{X}_e + \mathbb{\Lambda}\underline{X}_i \\ \tfrac{d\phi}{dt} - \nabla.\left(D_m\nabla\phi\right) = \tfrac{\phi_h(\underline{X}_i)-\phi}{\tau}.\end{aligned} \quad (31)$$

where $d/dt = \partial_t + v_s.\nabla$ is the material derivative with respect to the cellular flow. As such, the only effect of the cellular flow is to add a transport term to (6). The impact of $v_s$ on the stability properties of (6) presented in the main paper can then be investigated case by case either by modeling $v_s$ or by at least checking its order of magnitude. For example for the limb bud morphogenesis $v_s \approx 0.01\mu$m/s [58, 59], while from our estimates the typical velocity of the interstitial flow driven by spatial changes in the porosity $\sqrt{D_m/\tau} \approx 0.1 - 10\mu$m/s $\gg v_s$. We therefore do not expect that the account of cell rearrangements will substantially modify the thresholds of patterning instabilities given in the main paper as they are happening on slower timescale than our morphogen-driven pattern formation mechanism. A word of caution for this specific system though, as we note that our model requires the existence of an least transient elastic tissue structure (i.e. of the existence of some transient level of physical connections between the cells either directly via some adhesion or indirectly via the Extra-Cellular Matrix (ECM)). Such hypothesis and the measurements of the ensuing elastic moduli would need to be carefully scrutinized for this specific example.

A second simple modeling assumption that can be used to generalize our results is to suppose that morphogens are driving the cellular flow according to

$$v_s = \underline{\vartheta}.\nabla\underline{X}_e \quad (32)$$

where $\underline{\vartheta} = (\vartheta_A, \vartheta_I)$ is a vector of chemotactic sensitivities of the cell motion with respect to the morphogens concentrations. Such chemotactic role of morphogens has been demonstrated in particular examples such as shh in the drosophila [60]. In this case, (30) is a closed system of equations which stability can be investigated. Clearly, all the instabilities presented in the paper will remain valid for small enough $\underline{\vartheta}$. In the more general case, new instabilities associated with chemotaxis will appear. In particular, we can illustrate this in the simple case where only one morphogen is present as in Section 1.7.1 and in the limit considered in the main paper where $\gamma_A \gg \lambda_A \gg 1/\tau_A$ and $\tau/\tau_A \sim K_A \ll 1$. If that case, if $\chi_A = 0$, we have shown that no instability occurs in (18) as (20) is not satisfied. This is no longer true if we consider that $\vartheta_A > 0$ as we find the onset of an oscillatory (Hopf) instability when

$$\frac{(2\phi^*-1)\vartheta_A A^*}{D_m} < \frac{D}{D_m} < \frac{\tau}{K_A\tau_A} + \frac{\vartheta_A A^* \phi^*}{D_m} - 2\sqrt{\frac{(1-\phi^*)\vartheta_A A^* \tau}{D_m K_A \tau_A}}.$$

While the first inequality insures that chemotaxis is not strong enough such that it would make an infinite number of high modes unstable by surpassing Fickian diffusion, the second inequality shows that chemotaxis alone can be enough to destabilize the regularizing effects of both Fickian and hydrodynamic diffusion as well as the morphogen chemical turnover. In general, that type of instability will combine with the ones presented in the main text but a systematic investigation of such cross-effects is left to future works.

### 2 Historical background

In 1948, Turing's long time interest for biology took a new twist, largely influenced by the works of D'Arcy Thompson and Conrad Waddington [61]. In this context, he decided to study pattern formation and morphogenesis during embryonic development, developing computer programs to numerically solve the complex partial differential equations resulting of his abstraction of the biological question. In a letter dated of 12 February 1951, he wrote to a former colleague and friend, Mike Woodger [62]:

> "Our new machine is to start arriving on Monday. I am hoping as one of the first jobs to do something about 'chemical embryology'."

These "jobs" would be the foundations of Turing's famous paper "The Chemical Basis of Morphogenesis" [63]. In this article, he proposed an elegant mathematical model of self-organisation, where two reacting chemicals can spontaneously form periodic spatial patterns through an instability driven by their difference in diffusivity. This idea was not only very influencial amongst biologists such as J.M. Smith [64] or physicists such as I. Prigogine [65], but was a the stem of a new class of PDE models which became central in the development of modern mathematical biology [39, 66]. However, given the lack of molecular and cellular evidences available at that time, in a lot of studies, the emphasis was often placed on the striking similarity of the patterns obtained by the model with those observed in biological systems such as animal coat and skin pigmentation patterns, rather than on discussing the biological validity of the modeling assumptions. Quite astonishingly, in a letter to Alan Turing dated of 11 September 1952, Waddington already raised similar concerns about the applicability of Turing's reaction-diffusion model to biological developmental systems, questioning its limitation to reproduce some observed behaviours in embryonic development such as pattern scaling with tissue size or the generation of a spatial pattern of discrete cell types [67]:

> "I was extremely interested to read your recent paper on the chemical basis of morphogenesis. [...] I rather doubt, however, whether the kind of processes with which you were concerned play a very role in the fundamental morphogenesis which occurs in early stages of development. [...] Even in a case like the regeneration of tentacles in Hydra the final result seems to me more regular that one would expect from your type of mechanism. For instance, if one splits two hydras longitudinally and places them together so that the split edges heal and forms a single tube of double the normal diameter, this enlarged tube still regenerates the normal number of tentacles, although it would, of course, contain twice the normal 'chemical wave-lengths' around its circumference."

Turing was not naive about the limitations and underlying assumptions of his model and in fact already hints, in his article, at possible extensions to his theory considering the structure and mechanical properties of biological tissues [63]:

> "[...] one proceeds as with a physical theory and defines an entity called 'the state of the system'[...] The description of the state consists of two parts, the mechanical and the chemical [...] In determining the changes of state one should take into account: (i) The changes of position and velocity as given by Newton's laws of motion. (ii)The stresses as given by the elasticities and motions, also taking into account the osmotic pressures as given from the chemical data (iii) The chemical reactions (iv) The diffusion of the chemical substances. The region in which this diffusion is possible is given from the mechanical data.[...] The interdependence of the chemical and mechanical data adds enormously to the difficulty, and attention will therefore be confined, so far as is possible, to cases where these can be separated."

The original work of Hans Othmer on the transition between discrete and continuous models of pattern formation in tissues in the 1970's is maybe the closest example of this new research program tangentially set by Turing in his paper. In introduction of a review article entitled "Current problems in pattern formation" Othmer wrote [68]:

> "In Turing's work and virtually all subsequent work, the only mode of transport considered is diffusion. Furthermore, the model systems all deal with structureless, tightly coupled cells or their continuum analogs. Some future work should be directed toward: (a) the analysis of other modes of transport (b) more realistic models of cell and tissue structure and, (c) networks of cells that communicate only indirectly via the external medium."

As early attempts to fullfill this program, we would like to highlight the pioneering work of L.E. Scriven [69] who, by coupling mechanical stresses with a biochemical reactions to describe spontaneous fluid transport, paved the way to modern active gels theory and to the work of G.F. Oster [70], one of the father of mechanobiology.

### 3 Methods

#### 3.1 Numerical stability analysis

Linear stability analysis is performed numerically using conventional algebra packages in Mathematica.

#### 3.2 Numerical experiments

Numerical integration of models (10), (17) and (21) are performed in 1D with a custom made code written in Matlab and available on request to the authors. In brief, the code uses a mass conserving finite volume stencil in space and a first order implicit scheme in time.
